# Selecting Clustering Algorithms for IBD Mapping

**DOI:** 10.1101/2021.08.11.456036

**Authors:** Ruhollah Shemirani, Gillian M Belbin, Keith Burghardt, Kristina Lerman, Christy L Avery, Eimear E Kenny, Christopher R Gignoux, José Luis Ambite

## Abstract

**Background:** Groups of distantly related individuals who share a short segment of their genome identical-by-descent (IBD) can provide insights about rare traits and diseases in massive biobanks via a process called IBD mapping. Clustering algorithms play an important role in finding these groups. We set out to analyze the fitness of commonly used, fast and scalable clustering algorithms for IBD mapping applications. We designed a realistic benchmark for local IBD graphs and utilized it to compare clustering algorithms in terms of statistical power. We also investigated the effectiveness of common clustering metrics as replacements for statistical power.

**Results:** We simulated 3.4 million clusters across 850 experiments with varying cluster counts, false-positive, and false-negative rates. Infomap and Markov Clustering (MCL) community detection methods have high statistical power in most of the graphs, compared to greedy methods such as Louvain and Leiden. We demonstrate that standard clustering metrics, such as modularity, cannot predict statistical power of algorithms in IBD mapping applications, though they can help with simulating realistic benchmarks. We extend our findings to real datasets by analyzing 3 populations in the Population Architecture using Genomics and Epidemiology (PAGE) Study with 51,000 members and 2 million shared segments on Chromosome 1, resulting in the extraction of 39 million local IBD clusters across three different populations in PAGE. We used cluster properties derived in PAGE to increase the accuracy of our simulations and comparison.

**Conclusions:** Markov Clustering produces a 30% increase in statistical power compared to the current state-of-art approach, while reducing runtime by 3 orders of magnitude; making it computationally tractable in modern large-scale genetic datasets. We provide an efficient implementation to enable clustering at scale for IBD mapping and poplation-based linkage for various populations and scenarios.

## Background

Finding structure in networks, known as community detection, or clustering, is an important problem with a wide range of biomedical applications such as systems biology [1], population structure studies [2] and health information systems [3]. In the past decade, computational geneticists have found a new application for clustering algorithms in the context of Identity-By-Descent (IBD) mapping [4]. IBD mapping is an approach for rare variant association testing that leverages genotype data in the absence of directly observed variation for genomic discovery. This method relies on inferring haplotypes of the genome that have been co-inherited identically from a recent common ancestor and using them as the basis for association, under the assumption that these recent, shared haplotypes may co-harbour recently arisen rare variation not directly captured on genotyping arrays to be included within the association mapping framework.

Gusev et al.[4] found that, empirically, IBD mapping can yield up to forty times more statistical power than standard genome-wide association analyses (GWAS), specifically for rare genetic variants with less than 3% frequency in the population. Using IBD mapping, they found known and novel associations with case-control data gathered from a study in the United Kingdom. Browning et al.[5] also replicated a number of GWAS results through IBD mapping. Kenny et al.[6] used IBD mapping to fine-map known associations with plasma plant sterol levels in an isolated founder island population in Kosrae. Finally, Belbin et al.[7] utilized IBD mapping to identify the source of a common collagen disease in the Puerto Rican population of BioMe biobank.

IBD mapping has three main steps, as illustrated in Figure 1. In the first step, an IBD estimation algorithm finds shared genetic segments along a chromosome. The chromosome is divided into consecutive windows, and for each window a graph is generated where nodes represent the samples and edges represent IBD sharing in that window between the samples. We call these graphs local IBD graphs. In Figure 1, samples 3 and 5 share a segment IBD that covers window *n*+1, thus, in the graph generated for that window, they are connected by an edge. Local IBD graphs are usually comprised of multiple connected subgraphs, such as cluster 2 and cluster 3 in the window *n* + 1 graph.

**Figure 1.**
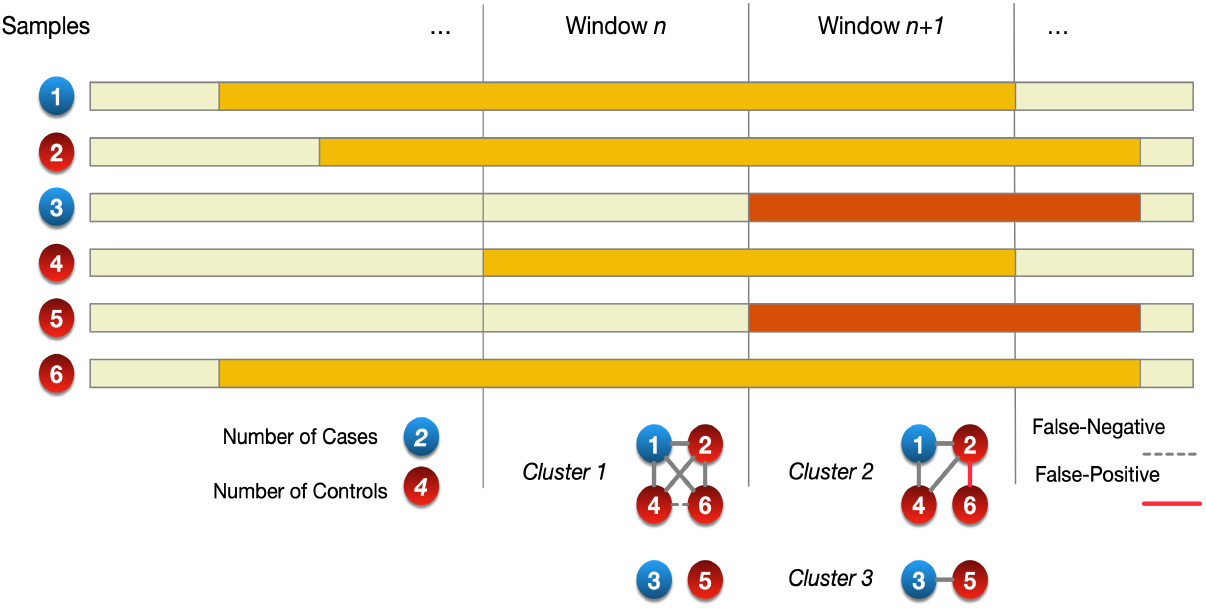
IBD Mapping Process. A general schema of the IBD mapping process. First, IBD segments are estimated and divided into windows. Second, a clustering algorithm finds the underlying communities in each window, eliminating false-positive and recovering false-negative edges. Third, a statistical test is conducted to find associations between cluster memberships and phenotypes. Significant dissimilarities between a family cluster and general population might be caused by an unrecorded rare variant carried on a shared IBD segment by the family.

In the second step of IBD mapping, a clustering algorithm extracts groups of related individuals in each local IBD graphs. In Figure 1, the clustering step finds 3 clusters. Clustering is necessary for the removal of false-positive edges (such as the edge between samples 2 and 6 on window *n* + 1) and the uncovering of missing false-negative edges (such as the edge connecting samples 4 and 6 on window *n*) in the local IBD graphs in order to accurately extract groups of distantly related individuals, who share a short segment of their genome IBD in the window. These segments might carry rare unrecorded causal genetic variants derived from a distant common ancestor that would otherwise not be detected using the genotype array data.

In the final step, each cluster is tested for associations with various traits and diseases. In Figure 1, while 1 out of 3 samples in the population are cases of the trait under study, in cluster 1, only one out of the four samples is a case, suggesting a lower susceptibility to the trait. Meanwhile, one out of the two samples in cluster 3 is a case, suggesting a higher susceptibility to the trait. Such associations with various traits might be caused by rare variants shared by members of the cluster.

While computational methods for the first step (IBD estimation), and the third step (statistical testing) of IBD mapping have been the subject of thorough studies [8, 9, 10], there has been a lack of advancement in IBD-clustering methodologies in the era of biobank-scale genomic data. There have been a plethora of innovations in clustering techniques in general however, due to their increased importance [11, 12, 13, 14, 15]. New clustering methods have been proposed to address the size of social networks and internet hosts, which have grown to many millions of nodes in the past decade[16, 17, 18]; or to find new community structures that reflect the underlying data more accurately[19, 20]. While the emergence of large biobanks necessitates the employment of such new clustering techniques in the context of IBD-mapping, it remains unclear how advancements in community detection methods translate to this process, where the unique population genetic properties of local IBD graphs may not resemble that of common graphs analyzed in other fields of study.

Graph properties are derived from factors such as i) the phenomena they represent, ii) the modeling approach used to generate them, and iii) the data collection methods used to ascertain them [21, 22]. Local IBD graphs have unique characteristics compared to graphs typically used in clustering method surveys. To explore this, we analyzed whether common graph properties such as connectivity, “small-world” property [22] and community size distributions [23] hold for local IBD graphs. Further, based on the properties of graphs being studied, definition and properties of clusters could also differ. Thus, in this manuscript, we also investigate the efficacy of common clustering metrics in the evaluation of clustering algorithms for IBD mapping purposes.

We design a realistic local IBD graph benchmark that addresses the shortcomings of common graph benchmarks in simulating local IBD graphs. We evaluate clustering methods based on their performance on simulated clusters that are generated to imitate real datasets. Furthermore, by simulating binary phenotypes for the synthesised clusters, our benchmark enables a comparison of clustering algorithms and metrics against each other in terms of their statistical power for the first time. In addition to finding an optimal approach for local IBD clustering, it also provided insight into the informativeness of common clustering metrics when the goal of community detection is to elucidate the underlying community and phenotype structure with high statistical power rather than analyzing the inherent structural properties of communities themselves.

## Methods

In this section, we first discuss the properties of local IBD graphs and compare them to common benchmark graphs. We demonstrate that common benchmarks do not realistically represent local IBD graphs. We then propose a new approach to simulate and benchmark local IBD graphs realistically. Secondly, we describe the clustering metrics we found relevant for IBD mapping along with our evaluation method for clustering algorithms. Finally, we describe the scalable clustering methods evaluated in this paper.

### Characterization of the Local IBD Graphs

Common benchmark graphs such as those introduced by Lancichinetti et al. [23], and Girvan and Newman [20], are used to evaluate clustering methods in a variety of fields [24]. However, they should not be used to simulate local IBD graphs, mainly due to the properties of the local IBD relationships that generates these graphs.

The topology of a graph that represents a relation *R* between entities of a set *S*, where an edge between two nodes *x, y* ∈ *S* means that (*x, y*) ∈ *R*, is dictated by the properties of the relation *R*. Local IBD relation is transitive, which means for every triplet of samples *a, b* and *c* in a dataset, if *a* and *b* share a segment *l*_1_ of their genome IBD, and *a* and *c* also share the same segment IBD, then *b* and *c* should also share *l*_1_ IBD. This means that, under ideal conditions, the local IBD relation can be represented as disjointed sets, making a graph representation inefficient, since it is solely made up of cliques. In practice, false-positive and false-negative edges obfuscate these cliques, necessitating a graph representation. The goal in the clustering of local IBD graphs is to recover these well-defined cliques.

Transitivity of local IBD relations results in uncommon graph properties. We look at the “small-world” property as an example [25]. This property measures the average distance between every two random nodes on a graph and compares it to a random Erdos Reney (ER) graph with the same size [22]. Small-world property cannot be calculated for local IBD graphs since, even before clustering, they are highly disconnected. To illustrate this, we analyzed the local IBD graphs of chromosome 1 in the “Population Architecture using Genomics and Epidemiology” (PAGE) dataset, which is a diverse genetic dataset with 52,273 samples from multiple populations [26]. We found that local IBD graphs in the PAGE dataset each have 13,961 connected components on average across 7952 local IBD graphs tested on chromosome 1; deriving an average of 3.74 nodes per connected component. Looking at two major subpopulations within PAGE, one enriched with IBD connections and recent founder effect, other with low levels of IBD sharing inside population, provides further evidence on disconnectedness of local IBD graphs. Puerto–Rican and African–American subpopulations in the PAGE dataset had 7.77 and 2.31 nodes per connected component respectively, suggesting that while population structure affects component size distributions, local IBD graphs are always highly disjointed. However, common benchmarking algorithms often generate a single connected component [23], in contrast with real world local IBD datasets. Cluster size distribution is another area of difference between local IBD graphs and others. The LFR benchmark [23], for example, only supports cluster size distributions that follow the power law. Estimating the local IBD cluster sizes using power law results in unrealistically low numbers for small clusters. For example, as demonstrated in Figure 2, a fitted power law distribution [27] underestimated the number of cluster sizes for clusters with less than thirteen members by a factor of ten. As discussed later in the results section, cluster size distribution affects the statistical power of Louvain and Leiden clustering algorithms [28]. Thus, while the tail distribution of local IBD graphs might follow the power law in some populations, using power law distributions (as is common in graph benchmarking [23]) for our simulations would result in an unrealistic evaluation the fitness of clustering algorithms to find local IBD communities since such distributions significantly underestimate the number of small clusters.

**Figure 2.**
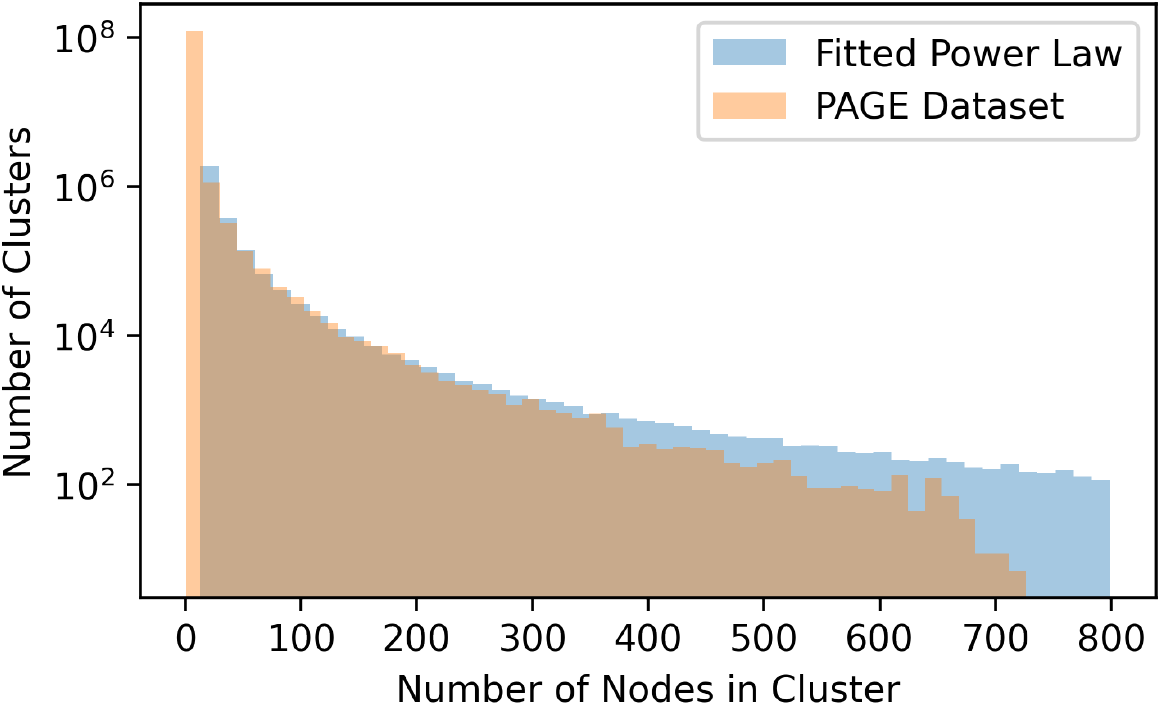
Cluster Size Distribution of Local IBD Clusters. The cluster size distribution of local IBD clusters along chromosome 1 in the PAGE study compared to a fitted power-law distribution. While the tail of the distribution can be approximated by the power-law, small clusters remain under-represented. The majority of local IBD clusters are small clusters. This causes the distribution to significantly diverge from the power-law approximations (p-value = 3 × 10^−14^).

Here, we describe a benchmarking approach that takes the specific properties of local IBD clusters into account. Our benchmark simulation has four steps. First, we generate a set of cliques, fully connected inside and disconnected from other cliques. These represent distinct IBD families in a given window. Members of a clique share the same DNA inherited from a common ancestor. To generate realistic clique size distributions, we sampled community size distributions on the chromosome one in the PAGE study dataset. In the second step, to simulate false-negative IBD segments, we randomly remove edges from each clique. Thirdly, in the false-positive edge generation step, we add inter-cluster edges to further increase the noise based on a given false-positive rate. Fourth, since the ultimate goal of IBD mapping is to test the family clusters for associations with traits, we simulate a set of binary traits, one for every cluster with more than 10 members. Traits have a significantly different prevalence among their designated cluster compared to the whole population. For example, trait *t*_*i*_, associated with cluster *ω*_*i*_, has a prevalence rate of *P*_*i*_ when every node (whole population) is tested. However, when only members of the cluster *ω*_*i*_ are tested, a different prevalence of 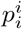 is achieved. The two prevalence rates *P*_*i*_ and 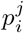 are chosen based on the cluster size and total node count such that the disparity between them is statistically significant (p-value_*binomial*_ ∼ 10^−10^). The significance threshold was chosen empirically.

### Metrics

Clustering algorithms categorize nodes into groups called clusters (The words cluster, community, class and category are often used interchangeably in clustering literature). Metrics help analyze various properties of the resulting clusters that are either related to the inherent features of the clusters, such as the density of connections in the clusters, or their concordance with the true structure of the graph, such as the number of nodes that are in the same clusters as they are in the ground truth. We call the first group feature-based metrics in this manuscript to distinguish them from metrics that are based on ground truth. For local IBD clustering, it is important to calculate how much the results reflect the true structure of the cliques underneath the noise and errors. We studied four metrics that aim to quantify such correlation with the ground truth through measuring information recovery. Since ground truth is often not available for real datasets, we also analyzed six clustering feature-based metrics to evaluate their efficiency in the absence of a ground truth.

#### Purity

Purity measures the the degree to which the clustering results replicate the ground truth clusters using a simple greedy mapping of the two [29]. To calculate purity, for every cluster *ϕ*_*j*_ recovered by the algorithm, we find a ground truth cluster *ω*_*i*_ with the highest number of common nodes and assign all the nodes in *ϕ*_*j*_ to *ω*_*i*_. Purity is then defined as the fraction of nodes that receive the correct cluster label using this approach. It is calculated using the following formula:

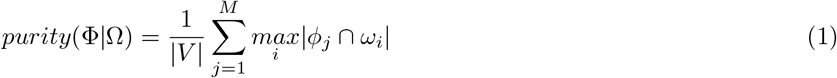

Where *V* is the set of nodes in the graph, Ω = {*ω*_1_, *ω*_2_, …, *ω*_*N*_} is the set of ground truth clusters, and Φ = {*ϕ*_1_, *ϕ*_2_, …, *ϕ*_*M*_} is the set of clusters extracted by the community detection algorithm. Purity is a simple metric to define and to calculate. It helps measure, transparently, how added noise can affect the structure of the clusters extracted compared to the original cliques. However, purity score can be maximized by assigning every node to a one node cluster of its own. For example, breaking down a cluster into two (or more) does not have an effect on the purity score, while it negatively affects power. Thus, for local IBD communities, where the majority of the clusters are doubletons, or tripletons, it can be misleading.

#### Normalized Mutual Information

Normalized Mutual Information (NMI) is built upon Shanon’s information theory [30, 31]. It measures the amount of information shared between the ground truth and the clustering results. It is calculated using the following formula:

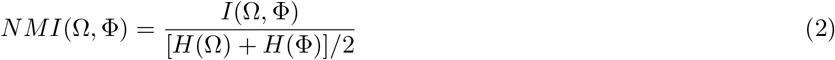

Where

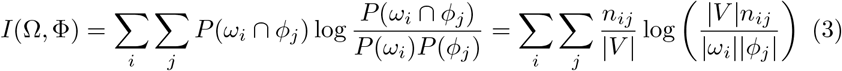

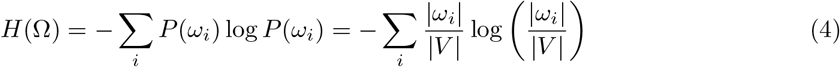

With |*V* | as the total number of nodes, *n*_*ij*_ as the number of nodes in *ω*_*i*_ ∩ *ϕ*_*j*_, and *H*(Φ) calculated using the same approach as *H*(Ω) (the entropy of the clustering and the entropy of the ground truth). Unlike purity, the value of NMI can only be maximized by replicating the same node assignments as the ground truth. The value of this score is 1 when the mutual information between Ω (ground truth) and Φ (clustering results) is maximized [31]. Compared to purity, increasing the number of clusters will not result in a perfect NMI score of 1, which makes it more dependable as a metric.

#### Adjusted Mutual Information

The baseline score in NMI can be improved through increasing the number of clusters. Vinh et al [32] first reported that the average NMI score of a random clustering increases as the number of clusters, or the size of the graph increases. To address this issue, they proposed the Adjusted Mutual Information (AMI). AMI is calculated by subtracting the expected mutual information (NMI score) of a random clustering of nodes from the NMI score of the real clustering. The random clustering should have the same number of clusters and number of nodes in each cluster.

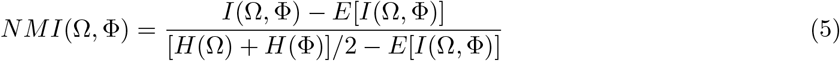

Where

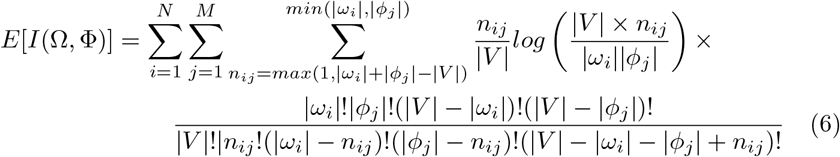

with all parameters calculated the same as NMI. Unlike NMI, a random assignment of nodes into clusters will yield an AMI score of zero. We report both NMI and AMI because the validity of the assumptions of the random assignment model of AMI is not universally accepted [33]. However, AMI is better calibrated. We confirmed that the AMI score has a zero baseline by randomizing the clustering results from Infomap across experiments so that the number of clusters and their sizes stays the same while the samples are shuffled among them. Figure 3A illustrates both NMI and AMI scores of the randomized clustering results. Randomized clusters continuously yielded a mean AMI score of zero and a mean NMI score higher than 0.6. The mean NMI score increases with the number of clusters. Adjusting the score to calculate AMI solves this problem.

**Figure 3.**
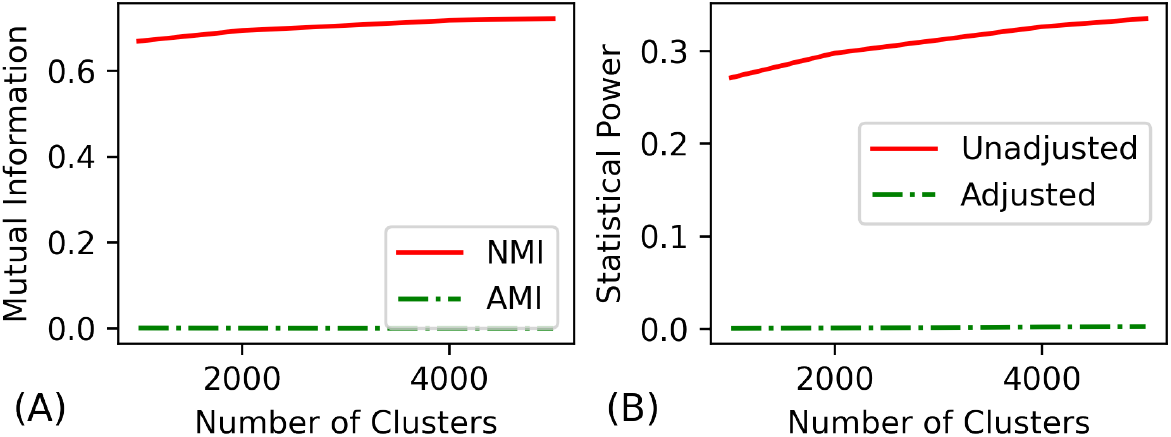
The effects of score adjustments on the baseline score. (A) Mutual information score (NMI), and (B) statistical power score of a random assignment of nodes to clusters has a non-zero baseline, which increases with the number of clusters. Adjusting these scores results in a score of zero for the same clustering. Adjustment is done through (A) subtracting the expected mutual information from NMI (AMI score), and (B) only including the results that pass the significance threshold when calculating power score. The scores were calculated by generating a random assignment of nodes to the same number of clusters and cluster sizes found by Infomap for each of the 750 simulations done in the paper.

#### Statistical Power

We simulate a binary trait *t*_*i*_ for every cluster *ω*_*i*_ that has more than 10 members. The prevalence 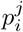 of the trait *t*_*i*_ in cluster *ω*_*i*_ is significantly different than the prevalence *P*_*i*_ of the trait among all nodes (p-value∼ 10^−10^). Statistical power measures the degree to which clustering algorithms can preserve this difference. It is calculated in 3 steps. First, for any extracted cluster *ϕ*_*j*_, the prevalence of every simulated trait *t*_*i*_ in that cluster, 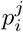, is calculated. Second, for each trait *t*_*i*_, the null hypothesis that the members of *ϕ*_*j*_ are as susceptible to test positive for *t*_*i*_ compared to other nodes in the graph (i.e., the difference between 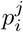 and *P*_*i*_ is statistically insignificant) is tested using a binomial test. The negated logarithm of the null hypothesis probability (p-value) is recorded as *S*(*ϕ*_*j*_, *t*_*i*_) only if it passes the bonferroni significance threshold in 5,000 experiments (p-value*<* 10^−5^). Third, for every cluster *ϕ*_*j*_, we only consider the largest score (*S*(*ϕ*_*j*_, *t*_*i*_)) value. Statistical power is then defined as the fraction of the maximum score (ground truth clustering score) retrieved by the clustering method. It is calculated using the following formula:

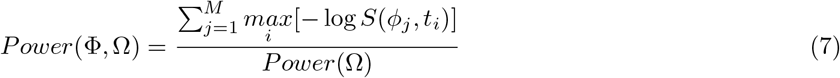

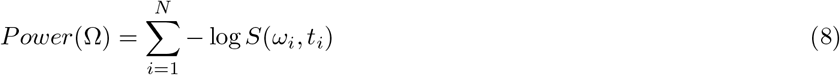

Statistical power ranges from zero to one. Random assignment of nodes to clusters results in a score close to zero; since the difference between the prevalence of any disease in clusters compared to the whole dataset becomes negligible. A clustering that exactly follows the ground truth results in a perfect score of one. Moreover, the value of this score cannot be optimized by extracting more fine-grained clusters as test scores that fail to pass the significance threshold are discarded, addressing the granularity problems with the purity and NMI scores. Figure 3B illustrates the effect of the significance threshold on the baseline power score using the same experiment we did for NMI/AMI comparison. Similar to NMI, a lack of correction for random results will increase the baseline statistical power score.

#### Modularity

Modularity measures the strength of clustering in terms of the density of links inside the clusters compared to external links connecting them [34]. It is desirable to have clusters densely connected within and sparsely connected to others [24]. Across all clusters, modularity measures the expected probability that any random link is located inside a cluster. It is calculated using the following formula:

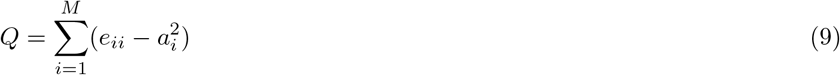

Where *M* is the number of clusters, *e*_*ii*_ is the fraction of edges that have both of their ends in the cluster *i*, and *a*_*i*_ is the percentage of edges that have at least one the their ends in the cluster *i*. Since modularity only measures the strength of a clustering, its calculation does not require ground truth information, in contrast with the previous metrics. We refer to the metrics that do not require ground truth information as feature-based metrics. Examination of modularity is essential since two of the algorithms analyzed in this paper use modularity optimization.

#### Other Feature-Based Metrics

In addition to modularity we analyze five other feature-based metrics. While modularity aims to quantify the overall strength of clusters, in terms of both their inner edge density, and outgoing connection sparsity, each of the following metrics only focus on one of the two.

- **Connectivity:** The percentage of nodes that are connected to more than half of other nodes in their cluster. This score offers an indirect way to measure the strength of the clustering. Being connected to more than half of a cluster’s members is impossible for the nodes in a “cluster” that is made up of smaller clusters with minimal false-positive connections.
- **Coverage:** The percentage of intra-cluster edges out of all edges [35]. Coverage captures an estimation of the fraction of edges that are deemed true-positive by the clustering algorithm.
- **Inter-Cluster Edge Rate:** The percentage of all edges that connect different clusters. This metric can be used to infer about the percentage of available edges that should be false-positives for the clustering to be correct. Subtracting coverage score from its maximum of 1 (where all edges are covered by clustering) yields this score.
- **Missing Intra-Cluster Edge Rate:** The percentage of the edges that are missing from clusters, assuming they are cliques, compared to the total number of edges expected in a set of cliques with the same number of members. This metric can help illustrate the rate of false-negatives necessary for the clustering to be correct.
- **Highly Connected Rate:** Out of all clusters that have more than ten members, the percentage of those that are highly connected. We define highly connected clusters as the ones that posses more than half of the edges of a clique with the same number of nodes. This score is similar to connectivity, however it only concerns graph sizes that are important to statistical power calculations. Also it has a binary nature in that either all the nodes in the cluster pass the test or none of them will.

### Clustering Methods

In this section, we describe the clustering algorithms evaluated in this study. We have chosen five algorithms in three categories based on their methodology. Every tested algorithm, except for Highly Connected Subgraphs(HCS), is scalable to large datasets [31], and can analyze our largest simulated dataset with 11,000 clusters in less than 5 minutes on average on our workstation running CentOS Linux release 7.4.1708 with 128 GB of memory and Intel® Xeon® Processors E5-2695 v2 (2.4 GHz) on a single thread.

#### Min-Cut Based Method

Min-cut based approaches use consecutive min-cuts to iteratively divide a graph into subgraphs that are highly connected inside and have few connections to other subgraphs [36]. The Highly Connected Subgraphs (HCS) algorithm is a example of a min-cut based approach. The criteria to decide whether a subgraph is highly connected depends on the algorithm [37]. A simple example is defining highly connected subgraphs as the ones where each node is connected to at least half of the other nodes in the subgraph. Whenever a subgraph reaches this threshold, the algorithm considers it a cluster. DASH [4], the best known local IBD clustering tool, uses a modified version of HCS. To decide whether a subgraph is highly connected or not, DASH uses an optimization function which requires parameter tuning using *a priori* estimation of the rate false-negative and false-positive edges in the graph. To reduce the complexity of our experiments, and to have a fair comparison, we use the fast HCS algorithm described in [38], where a subgraph is called highly connected if a minimum of |*V* |*/*2 edges need to be removed to break it down into two disjointed subgraphs. Running HCS is equivalent to running DASH without passing the false-positive/false-negative information to the algorithm.

#### Modularity Optimizing Approaches

Modularity optimization approaches aim to maximize the modularity score function described in the metrics section[34]. Modularity optimization can uncover structures unknown *a priori* [39]. Moreover, high modularity is a prefered structural property for clusters as it indicates the strength of internal connections compared to external ones [40]. While optimizing modularity is NP-complete [40], greedy algorithms have proven to be fast and approximately accurate [39, 41, 24, 31]. Utilizing greedy heuristics, methods such as the Louvain algorithm achieve a polynomial runtime [39]. We also analyze the Leiden algorithm [41], another clustering algorithm based on greedy optimization of modularity. In Leiden algorithm, Traag et al. have improved Louvain clustering heuristically in scenarios that cause the Louvain algorithm to find clusters that are poorly-connected, or entirely disconnected [41].

#### Information Theory Based Methods

Instead of focusing on the hierarchical structure of clusters and their connectivity, methods based on information theory focus on the flow of information among the nodes. These methods often analyze information flow using random walk probabilities between the nodes. We analyze two methods in this category.

#### Infomap

The Infomap algorithm is the most commonly used information theory based clustering method. It aims to minimize the memory required to describe any random walk on the graph, also called path description length, through collapsing groups of vertices that are more likely to appear consecutively in a random walk into a single vertex. These groups are then labeled as clusters. Infomap uses an objective function called the Map Equation to approximately minimize the path description length [42]. This approximation approach helps infomap to scale to large graphs [31]. For every pair of nodes connected by an edge, Infomap checks whether collapsing them into a single node would help minimize the result of the map equation (and by definition, the path description length) or not; if so, it puts them in the same cluster.

#### Markov Clustering

The Markov Clustering (MCL) algorithm uses flow dynamics to simulate random walks on the graph until they converge to a steady state [43]. To do so, it employs matrix multiplication to simulate the destination probabilities of a single step of a random walk and then inflates those probabilities to weaken the unstable paths in an effort to guarantee that the series of random walk simulations will always converge to a specific, non-random group of clusters. The elimination rate of the lower-probability paths is controlled through a parameter called the inflation rate with a range of 1.2 to 5.0. Theoretically, setting a higher inflation rate will result in finer-grained clusters with dense connections. We explore four inflation rates: 1.5, 2, 3, 5 in the results section. We use subscripts to denote which parameter was used to run MCL: *MCL*_1.5_, *MCL*_2_, *MCL*_3_, *MCL*_5_.

## Results

### Performance on Simulated Data

Using our benchmark algorithm described in the methods section, we generated 750 graphs (one per experiment). The number of clusters in each experiment ranged from 1,000 to 5,000. For each cluster count, we considered 25 sets of false-positive and false-negative combinations ranging from 5% to 50%. Finally, for each combination of cluster count, false-positive, and false-negative rates we simulated six experiments. The mean and standard error of the prevalence of simulated traits among the whole population in each of these six experiments were selected from a set of 3 (0.05, 0.1, 0.15) and 2 (0.01, 0.1) predetermined values, respectively; for a total of 750 experiments. This added up to a total of 2,274,500 clusters with more than 6 million nodes across all simulated experiments. We demonstrate the accuracy of our benchmark in Figure 4. The figure illustrates the layout of a random sample of local IBD graphs on chromosome 1 (Figure 4 A) against randomly generated benchmarks, both using our algorithm (Figure 4 B) and the LFR algorithm (Figure 4 C). As shown in the figure, our benchmark simulates the disjointedness of local IBD graphs, unlike the LFR algorithm.

**Figure 4.**
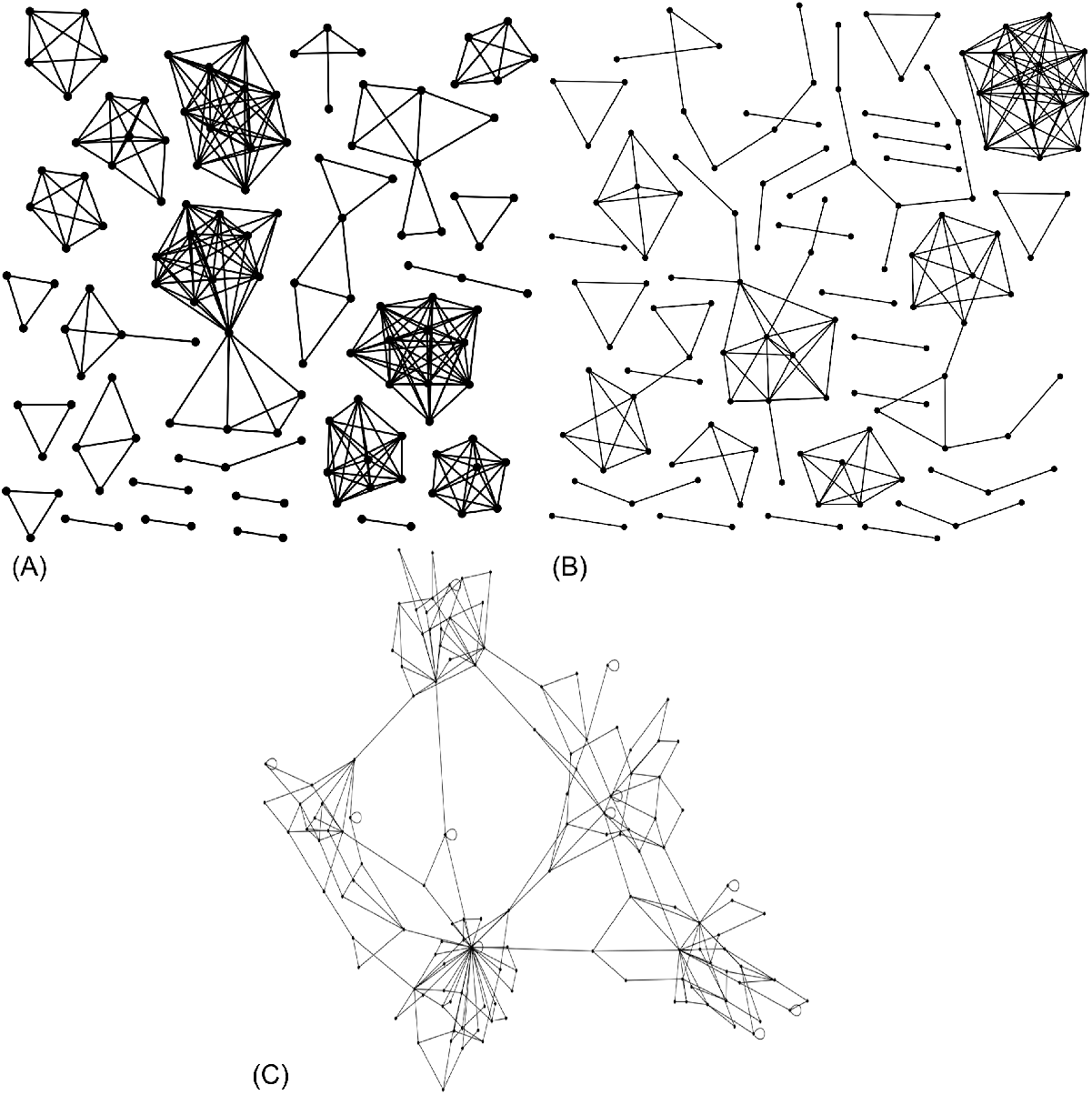
Comparison of graph layouts. Graph layouts with ∼ 130 nodes generated (A) from the real IBD data in the PAGE study dataset on chromosome one via random sampling, (B) by our benchmark algorithm, (C) using the LFR method. LFR benchmarks are generally comprised of a single connected component which does not happen in real local IBD graphs. Further, the cluster sizes generated using the power law distribution do not resemble that of the real local IBD graphs. Our benchmark addresses both of these issues.

We ran the clustering algorithms on these simulated datasets to extract clusters. We then calculated the scores achieved by every method for each metric described earlier. In the remainder of this section, we first discuss the connections between the clustering metrics. We then describe the performance of the clustering algorithms on simulated data using the metrics. Finally, we discuss how their performance on simulated data can be translated to real datasets.

#### Clustering Metrics

We calculated the Pearson correlation coefficients and *R*^2^ scores [44] between metrics across all simulations for every tool to see whether, and to what degree, each clustering metric is associated with statistical power. The results are displayed in Figure 5.

**Figure 5.**
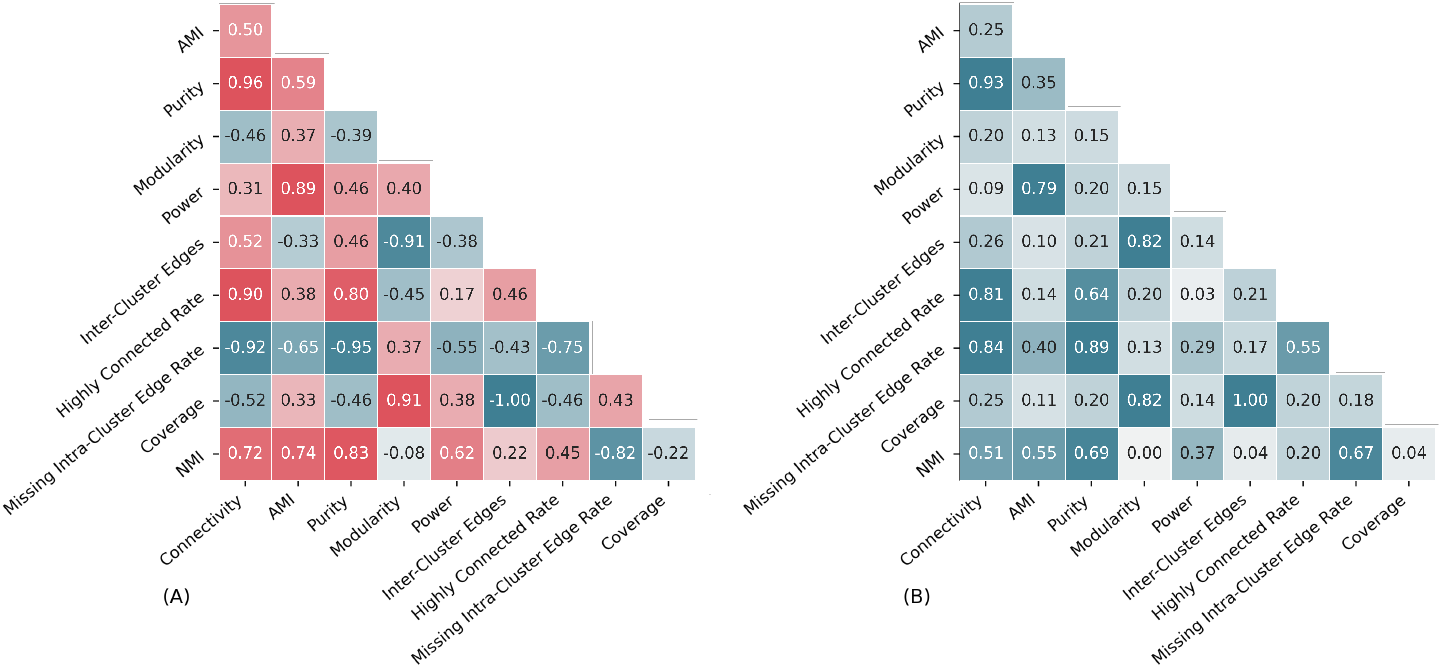
Similarities Among Clustering Metrics. (A) Correlation, and (B) *R*^2^ scores among clustering metrics across all simulations. AMI has the highest correlation and *R*^2^ with statistical power score. Among feature-based metrics, missing intra-cluster edge rate can predict power better than the others.

Among all metrics, AMI had the highest concordance with statistical power (Figure 5 B), explaining 79% of the variation of the power score. Among the feature-based metrics, missing intra-cluster edge rate has the highest *R*^2^ score, predicting 29% of the variability of statistical power, while highly connected rate had the lowest score. Thus, while generating denser subgraphs with less missing edges is important to gain power, focusing solely on the density and ignoring coverage will counter those effects, resulting in lower power.

Modularity showed a weak association with statistical power (*R*^2^ = 0.14). It was not able to predict statistical power as precisely as missing intra-cluster edge rate. Thus, partitioning a graph into highly modular subgraphs (through optimizing modularity) does not necessarily result in clusters that represent the true IBD communities in the underlying population. While optimizing modularity is advantageous in finding large non-clique-like communities [45], local IBD graphs are both clique-like and often smaller in scale (See Figure 2). A high percentage of small cliques results in amplification of discordance between modularity and power scores. Adding or removing a node from a cluster with a large enough number of members, say two hundred, to improve the modularity score does not drastically change the statistical power of the clustering. However, this is not the case for smaller clusters, for example, those with ten to twenty members.

To further demonstrate the effects of small clusters (prevalent in local IBD cluster size distributions) on the discordance between modularity and statistical power, we re-ran the same experiments with a uniform cluster size distribution (instead of our sampled distribution). The *R*^2^ score for modularity and statistical power rose to 0.34 (from 0.15) and the gap between modularity/power and AMI/power *R*^2^ scores decreased from 0.63 with realistic distribution, to 0.49 with the uniform distribution. AMI/power *R*^2^ score slightly increased to 0.83, compared to the 100% increase in modularity/power *R*^2^ score.

The observed discordance between modularity and power in our experiments can also be explained through the concept of “resolution-limit” in modularity optimization, i.e., the inability of modularity optimizing methods in detecting fine-grained clusters. Fortunato and Barthelemy [28] found that the modularity score for a clustering is not only dependant on the structure of the graph, but also on the expected maximum possible modularity of any random graph with the same number of edges. Through the introduction of the resolution limit, they further illustrated that modularity optimization may fail to capture clusters that have an order of magnitude fewer edges compared to the total number of edges in the graph. In other words, collapsing cliques with minimal connectivity to each other (two cliques connected via a single edge) and no connectivity to the rest of the graph could increase modularity of a clustering due to the resolution limit. Small, clique-like structure of local IBD graphs intensifies the effects of this phenomenon on the performance of the madularity metric and methods optimizing it; especially when compared to the leading methods/metric. We further discuss the resolution limit when analyzing the performace of the clustering algorithms.

Our results show that purity is unfit for our IBD clustering purposes. Specifically, regardless of the true underlying structure, a more granular clustering always yields a higher purity score. This is further demonstrated in Figures 6, 7 and 8, where we look at the score trends of all clustering algorithms in every metric across our simulated data as the number of clusters, false-negatives, and false-positives grow, respectively. Consider Figure 6 as an example. *MCL*_5_, a clustering approach that has the fifth best performance in statistical power (Figure 6 E), repeatedly gains the highest purity score (close to the perfect score of 1 in Figure 6 C), due to over-clustering, suggesting that purity score in the absence of others can be misleading and uninformative.

**Figure 6.**
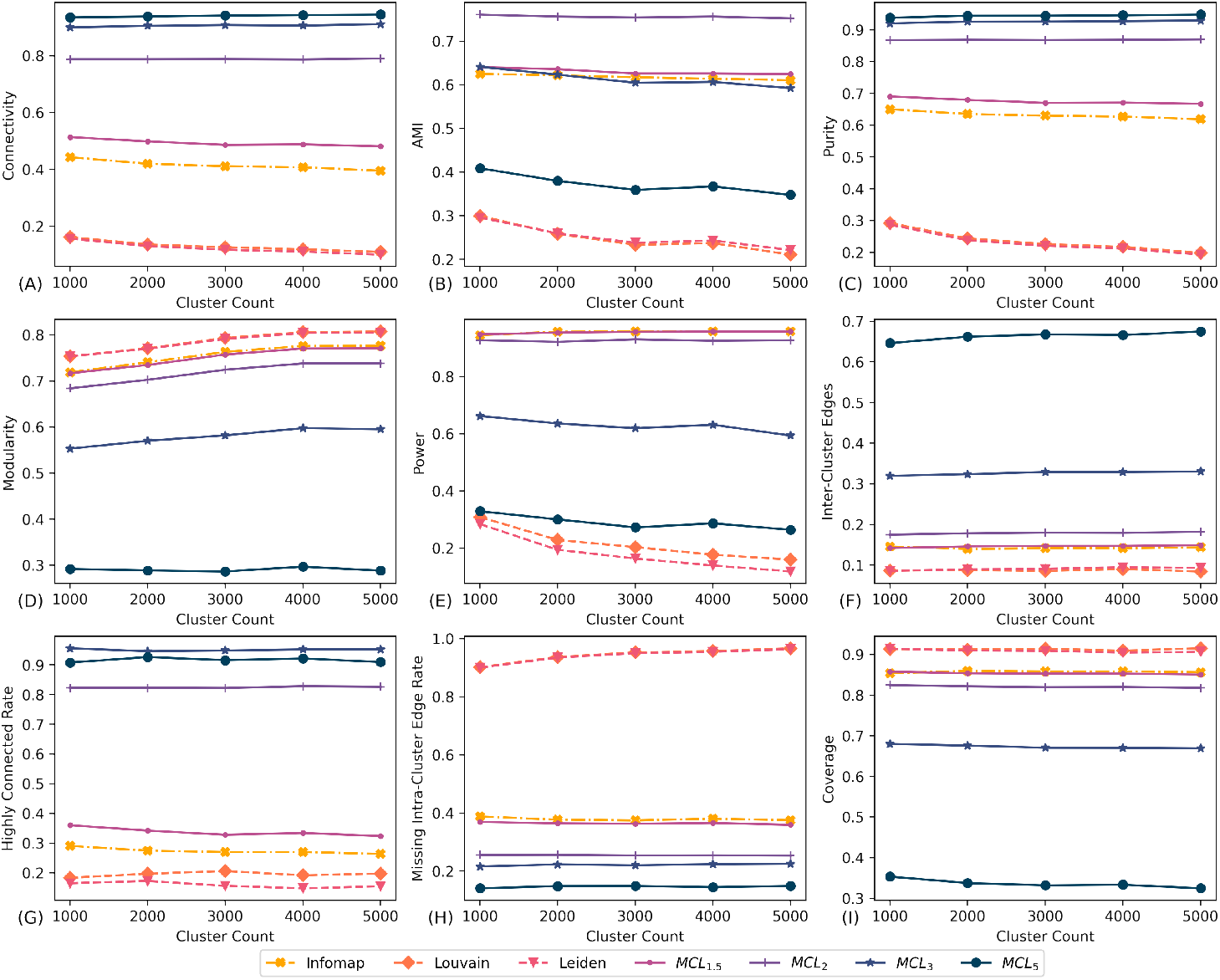
The effects of cluster count on the scores of algorithms. The effects of number of simulated clusters on the performance of algorithms through different metrics using 750 simulated graphs sampled from the PAGE dataset. With false-positive and false-negative edges ranging from 5%-50%. Total number of clusters affected the performance of some algorithms. Such dependency on the number of clusters negatively affects IBD mapping application.

**Figure 7.**
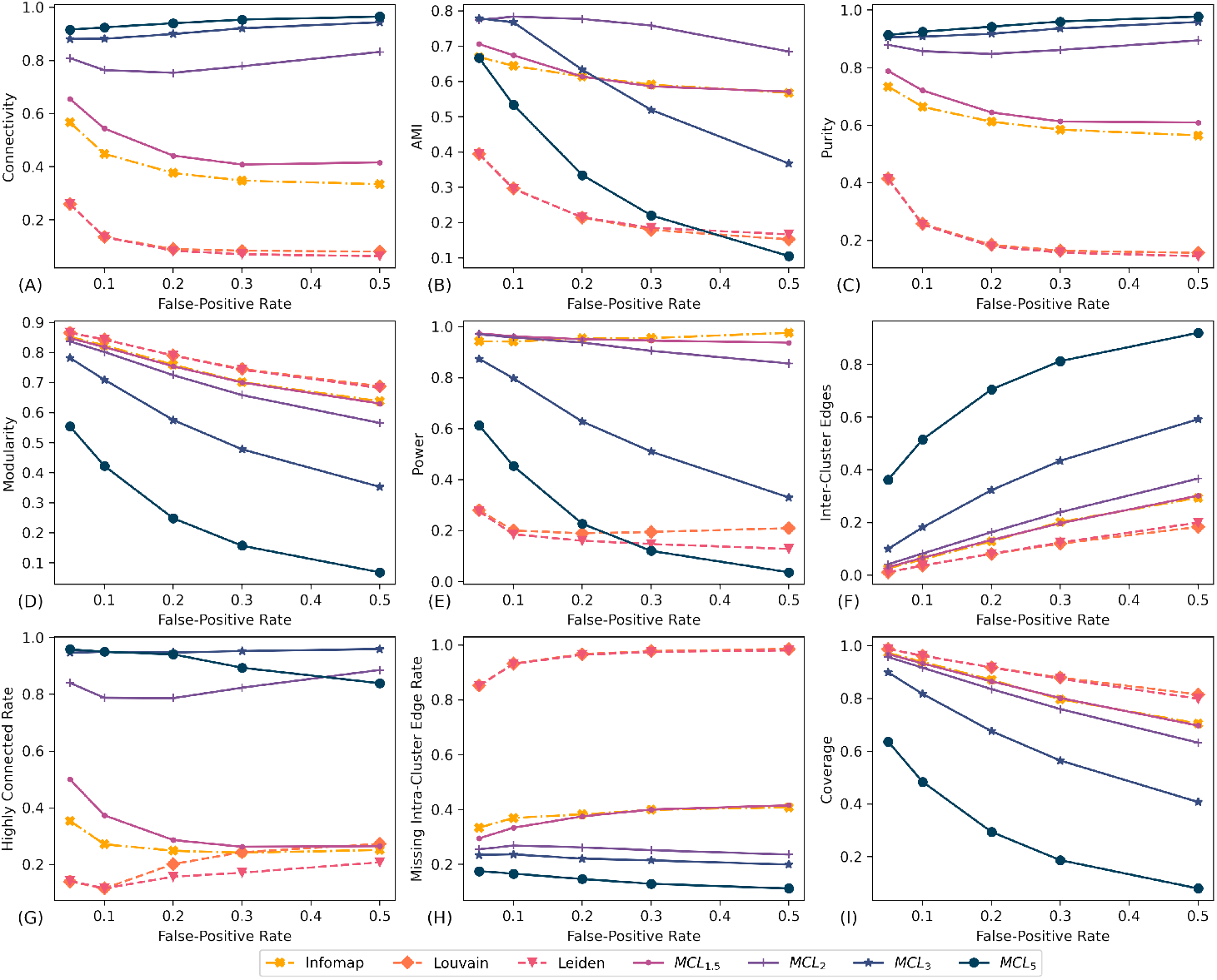
The effects of false-positives on the scores of algorithms. The effects of false-positive rate on the performance of algorithms through different metrics using 750 simulated graphs sampled from the PAGE dataset. False-negative edges ranging from 5%-50% and the number of simulated clusters per graph ranging from 1,000 to 5,000. Infomap and *MCL*_1.5_ had the most stable performances across various rates of false-positives, followed closely by *MCL*_2_. In real dataset, we expect the false-positive rate to never go above 15%.

**Figure 8.**
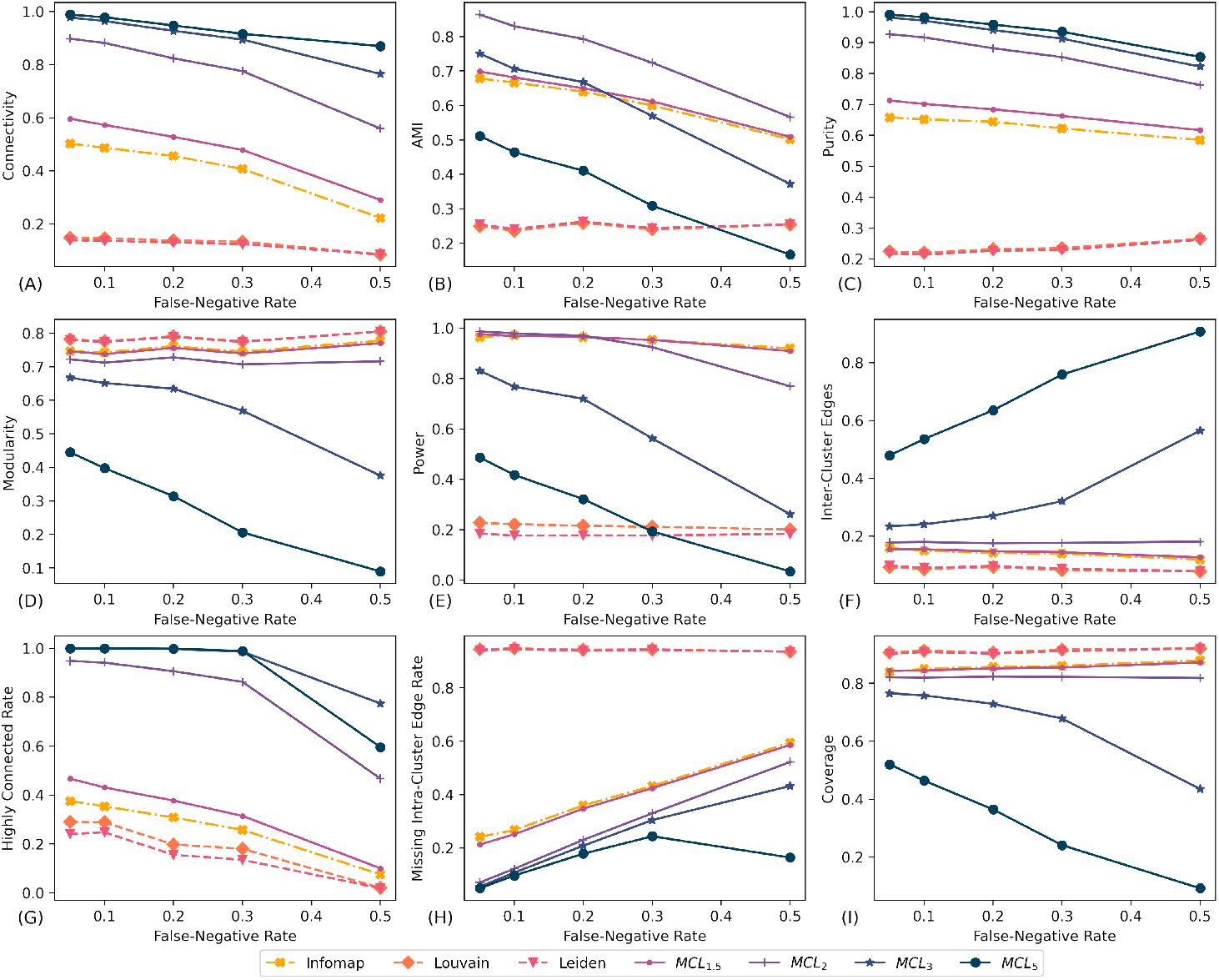
The effects of false-negatives on the scores of algorithms. The effects of false-negative rate on the performance of algorithms through different metrics using 6,250 simulated graphs sampled from the PAGE dataset. False-positive edges ranging from 5%-50% and the number of simulated clusters per graph ranging from 1,000 to 5,000. Compared to observations with the number of clusters and rate of false-positives, Louvain and Leiden have a stable performance when false-negative rate is increased. Although they are still under-performing against Infomap and MCL with an 80% statistical power score gap.

We have found the AMI score to be the best indicator of statistical power among the metrics we tested. However, due to the effects of smaller clusters (with less than 10 nodes) on AMI, its concordance with statistical power is imperfect. As further demonstrated by the performances of *MCL*_2_ and *MCL*_3_ in Figures 6, 7, and 8, compared to statistical power (Figure 6 E), the gap between *MCL*_3_ and top performing methods is less pronounced for the AMI scores (Figure 6 B). Moreover, *MCL*_2_ performance increases and surpasses the performance of Infomap and *MCL*_1.5_ in terms of AMI score compared to the statistical power. The same issue, together with a high baseline, severely affects the performance of NMI as well. Figure 9 illustrates the NMI score trends of every method across our simulated data as the number of clusters (A), false-negative edges (B), and false-positive edges (C) grows. Compared to AMI scores, the gap in the NMI scores of MCL algorithms and Infomap is even less pronounced, to a point where *MCL*_5_ and *MCL*_3_ receive the same average score as Infomap while statistical power score of Infomap is on average 57% and 27% higher than *MCL*_3_ and *MCL*_5_, respectively.

**Figure 9.**
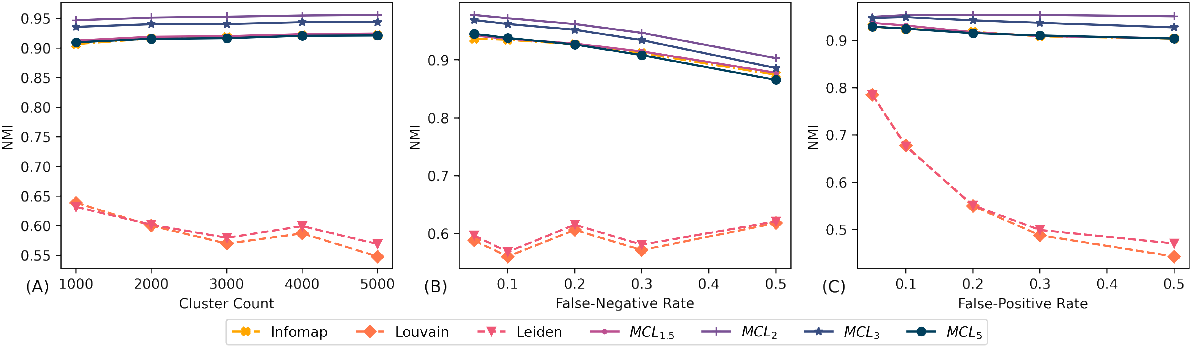
NMI score trends. The NMI score of algorithms as (A) the number of clusters, (B) false-negative rate, and (C) false-positive rate grows over all experiments. For MCL and Infomap, an increased number of clusters can increase NMI score due to a lack of adjustment for the probability of a random clustering conforming to the ground truth. Louvain and Leiden are the exceptions here as they undergo the effects of resolution-limit.

Another disadvantage of the AMI metric is its reliance on the existence of ground truth data. However, in the absence of the true clustering information, our experiments show that none of feature-based metrics, including modularity, can be used to accurately predict statistical power. We look at missing intra-cluster edge rate as an example due to its higher *R*^2^ score. Methods that yield the highest and lowest score in this metric (Leiden and *MCL*_5_) both perform poorly in terms of statistical power (see Figure 6). Thus, unlike AMI, this, and other feature-based metrics, do not preserve the algorithms rankings in terms of statistical power. We cannot rely upon these metrics as a representation of statistical power of a clustering.

#### Clustering Algorithms

Table 1 shows the average score of clustering algorithms for every metric across all of the simulated datasets. Infomap received the highest average statistical power score, followed closely by MCL, while Louvain and Leiden got the lowest score.

**Table 1.** Average scores (with standard error) of clustering algorithms across our experiments. Overall, MLC_2_, Informap, and MCL_1.5_ yielded the best performances. Modularity optimizing methods had a much lower power..

|  |  | Methods |  |  |  |  |  |  |
| --- | --- | --- | --- | --- | --- | --- | --- | --- |
| Metrics | | Infomap | Louvain | Leiden | $MCL_{1.5}$ | $MCL_2$ | $MCL_3$ | $MCL_5$ |
| Connectivity | Mean | 41.52% | 13.03% | 12.28% | 49.34% | 78.78% | 90.58% | **94.03%** |
|  | Error | 14.20 | 8.79 | 9.07 | 15.63 | 13.31 | 8.31 | 5.32 |
| AMI | Mean | 61.77% | 24.81% | 25.18% | 63.05% | **75.60%** | 61.36% | 37.26% |
|  | Error | 8.62 | 12.04 | 11.65 | 9.51 | 12.69 | 22.12 | 26.02 |
| Purity | Mean | 63.24% | 23.58% | 23.03% | 67.58% | 86.85% | 92.57% | **94.41%** |
|  | Error | 7.62 | 11.84 | 12.12 | 8.77 | 6.76 | 6.51 | 6.01 |
| Modularity | Mean | 75.52% | **78.64%** | 78.48% | 75.02% | 71.76% | 57.98% | 29.07% |
|  | Error | 13.52 | 11.99 | 12.17 | 13.19 | 14.32 | 21.14 | 24.39 |
| Power | Mean | **95.49%** | 21.54% | 17.98% | 95.47% | 92.61% | 62.85% | 29.05% |
|  | Error | 5.84 | 10.36 | 10.98 | 3.69 | 12.19 | 31.64 | 30.97 |
| ICE* | Mean | 14.26% | **8.70%** | 9.10% | 14.64% | 17.91% | 32.66% | 66.35% |
|  | Error | 10.61 | 6.77 | 7.45 | 9.80 | 11.66 | 21.84 | 27.19 |
| HCR** | Mean | 27.40% | 19.53% | 15.95% | 33.80% | 82.50% | **95.15%** | 91.67% |
|  | Error | 12.27 | 19.35 | 16.42 | 17.11 | 19.47 | 9.56 | 23.31 |
| MICER*** | Mean | 37.92% | 94.33% | 94.09% | 36.44% | 25.51% | 22.17% | **14.65%** |
|  | Error | 13.93 | 6.24 | 6.12 | 14.64 | 16.20 | 13.84 | 8.04 |
| Coverage | Mean | 85.74% | **91.30%** | 90.90% | 85.36% | 82.09% | 67.34% | 33.65% |
|  | Error | 10.61 | 6.77 | 7.45 | 9.80 | 11.66 | 21.84 | 27.19 |
| NMI | Mean | 91.72% | 58.89% | 59.65% | 92.02% | **95.26%** | 94.10% | 91.70% |
|  | Error | 3.58 | 19.47 | 19.19 | 2.90 | 2.89 | 3.36 | 3.58 |

\*: Inter-Cluster Edges \*\*:Highly Connected Rate \*\*\*: Missing Intra-Cluster Edges Rate

As expected, Louvain and Leiden algorithms yield the most modular clustering results; followed by Infomap. In terms of conforming to the ground truth (purity, power, and AMI/NMI scores), however, Louvain and Leiden achieve a much lower score than MCL and Infomap. Since Louvain and Leiden algorithms are both based on optimizing modularity, their performance further corroborates our analysis of resolution limit in the previous section.

As a result of resolution limit, Louvain and Leiden were unable to find smaller communities in our simulations. We know from [28] that greedy modularity optimization tends to merge lightly connected subgraphs into clusters. Although clusters of any size can be incorrectly merged when optimizing modularity, the most affected clusters are those with fewer internal edges than 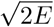, with *E* as the total number of edges in the graph. For example, the average number of edges for a graph with 2,000 clusters in our experiments is 62,007, which means any pairs of clusters that have a combined edge count smaller than 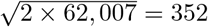 have a high chance of being merged by Louvain and Leiden if they are connected by a single edge, as it increases the modularity score. The vast majority of IBD clusters in our experiments have less than 352 individuals.

Figure 10 shows how the threshold for the smallest community size detectable by Louvain and Leiden is affected as the number of (A) clusters, (B) false-negative edges, and (C) false-positive edges, increase in our experiments. As illustrated in Figure 10A, the threshold for resolution limit grows at a faster rate compared to the number of large clusters. In other words, in local IBD graphs, the average number of subgraphs that are larger than this threshold decreases as the total number of clusters increases. The approximate threshold for resolution limit grew from 227 to 744 as cluster count was increased from 1,000 to 10,000 clusters. At the same time, the percentage of clusters larger than the resolution limit threshold decreased from 23.4% to 0.8%. As the size of the local IBD graphs grows, resolution limit causes Louvain and Leiden algorithms to merge subgraphs with minimal connections to each other, going as far as grouping sets of cliques connected by a single false-positive edge. This effect also causes the modularity optimizing algorithms to have an improved modularity score as the number of clusters grows while their statistical power decreases as shown in Figure 6D and E, respectively.

**Figure 10.**
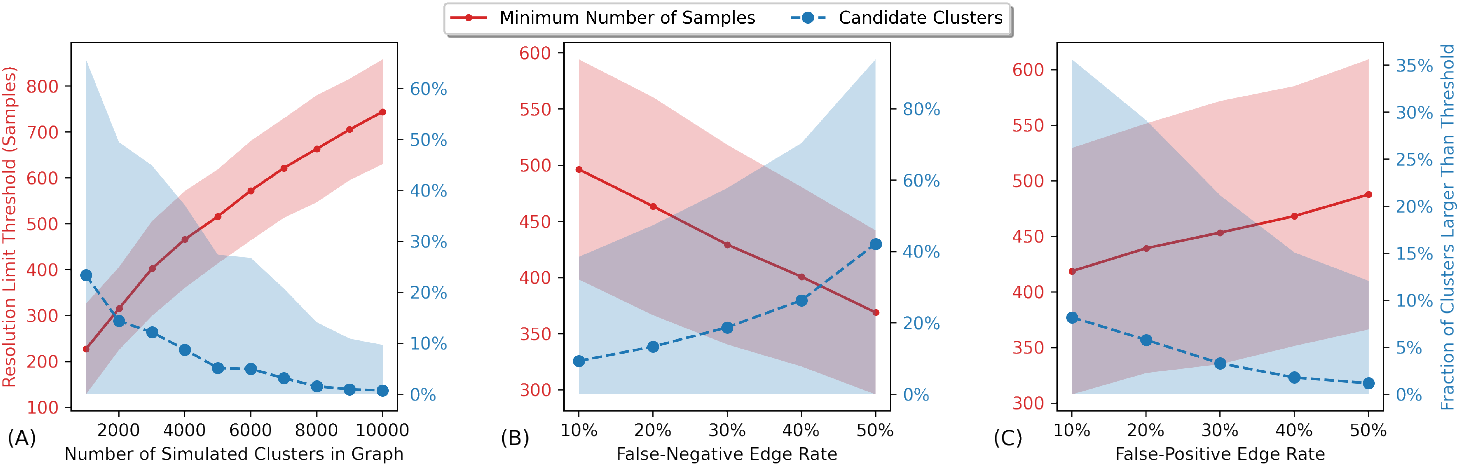
Resolution-limit trends. Trends for the average resolution limit threshold (red), and the expected frequency of clusters that pass the resolution limit threshold (blue) as (A) the number clusters grows, (B) number of false-positive edges grows, and (C) number of false-negative edges grows in 100 simulations sampled from the PAGE study dataset. Clusters that are not large enough to pass this threshold may be merged with other clusters in modularity optimization clustering process. The shaded area shows 95% confidence interval.

We further analyze the distribution of connectivity scores achieved by the algorithms across all of our simulations in Figure 11. The average percentage of nodes that were connected to at least half of the other members of their cluster, extracted by Louvain and Leiden, was 13% and 12%, respectively. The same average for *MCL*_2_ was 78%, indicating that Louvain and Leiden merge more cliques together compared to other methods.

**Figure 11.**
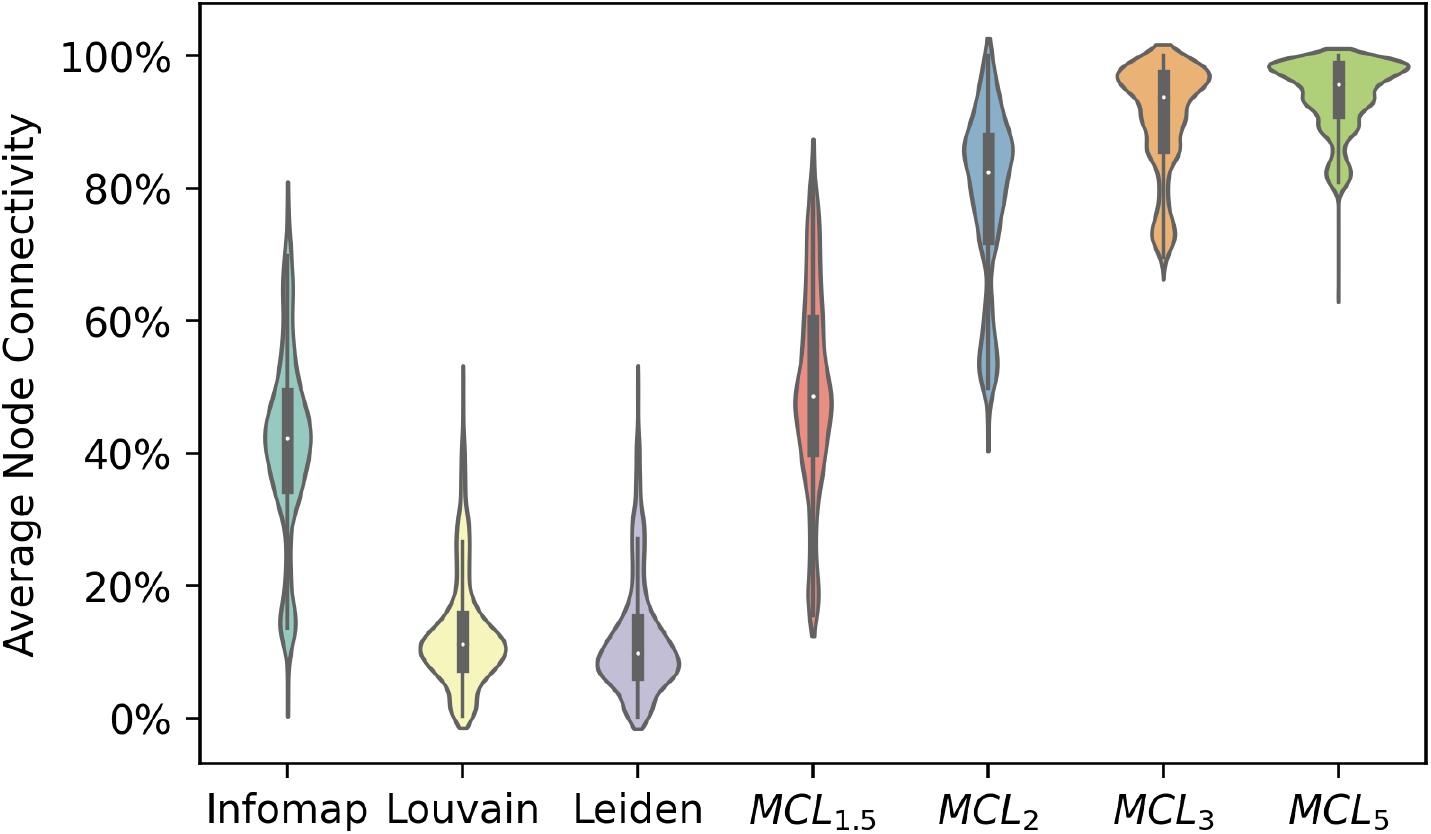
The effect of resolution-limit on the density of clusters. A violin plot for average connectivity score of clustering algorithms across our 750 simulations. Among the top performing algorithms in terms of power, *MCL*_2_ had the highest connectivity score at 79%, suggesting that it finds densely connected clusters that also conform to the ground truth. Infomap and *MCL*1.5 gained mean connectivity scores of 41% and 49% respectively. The lowest average scores belonged to Louvain and Leiden at 13% and 12%, respectively. Connectivity scores of Louvain and Leiden further demonstrate the inability of Louvain and Leiden to accurately extract clusters due to the resolution limit.

Resolution limit has another disadvantage; its accuracy depends on the overall edge count and not on the individual clusters [28], making it problematic for local IBD clustering; where a variety of cluster size distributions exist for the same total edge count. For example, in the PAGE study dataset, the average number of edges per cluster for local IBD graphs that only include samples from Puerto Rican and African American populations is 96.8±12.7 and 1.6±0.1 respectively. Thus, the statistical power of Louvain and Leiden is subject to change between the two populations, even if they are in the same dataset.

Figure 12 displays the distribution of the average number of nodes per cluster across 100 experiments each with 2,000 simulated clusters and false-positive/false-negative rates of 25%. The figure shows the tendencies of *MCL*_3_ and *MCL*_5_ for over-clustering and that of Louvain and Leiden for under-clustering that results in very low (for *MCL*_3_ and *MCL*_5_), and very high (for Louvain and Leiden) average number of nodes per clusters compared to the ground truth. While the average number of nodes per cluster in the ground truth was 3.6 (std=0.2), the average number of nodes per clusters found by Louvain and *MCL*_5_ were 197.7 (std=212.5) and 3.0 (std=1.7), respectively.

**Figure 12.**
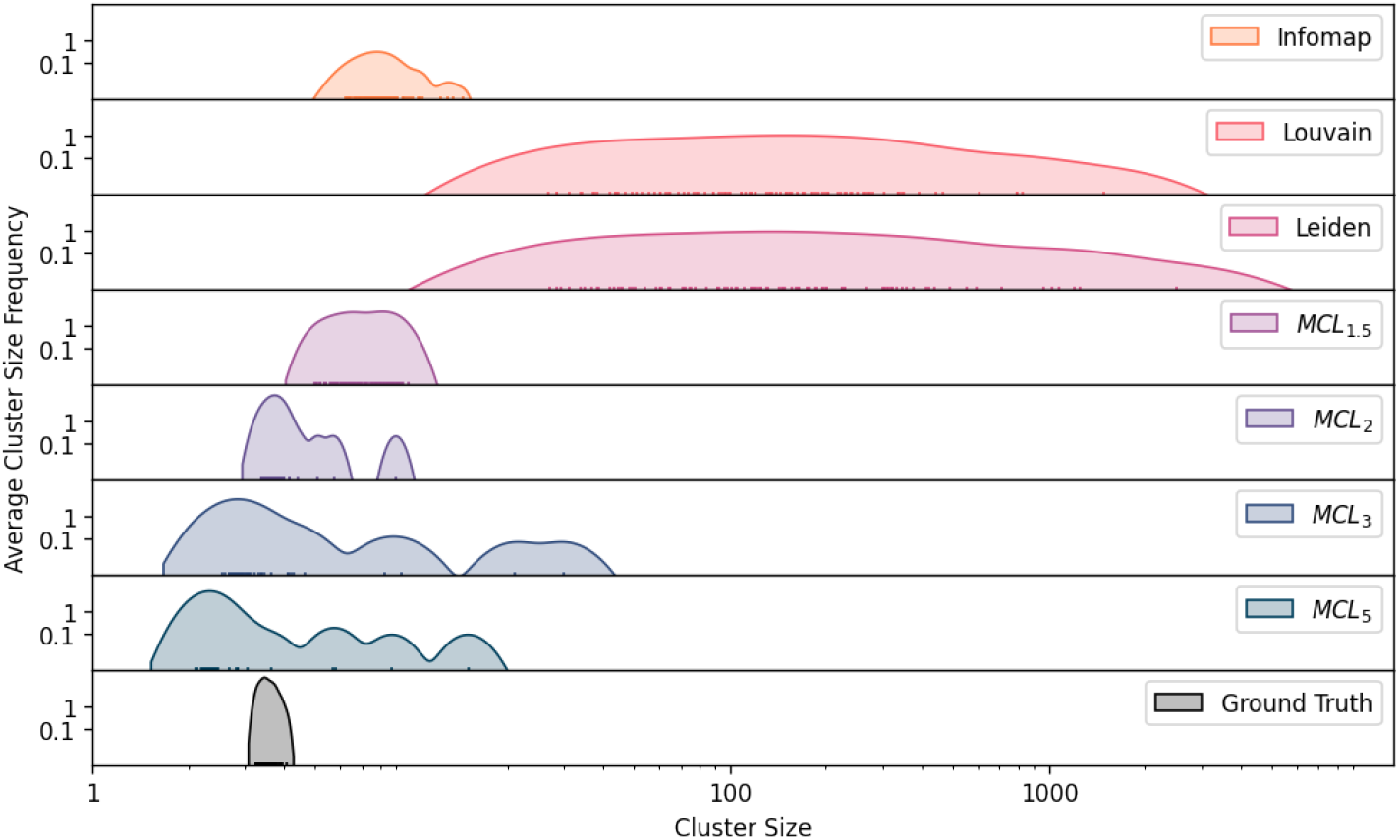
Extracted cluster size distribution. Distribution of the average number of nodes per cluster found by each algorithm across 100 simulations. Each simulated graph was comprised of 2,000 clusters, sampled from chromosome 1 in the PAGE study dataset with 25% false-positive and false-negative edges added. *MCL*_2_ had the closest distribution to the ground truth, while modularity optimizing methods had the furthest distribution. They failed to capture many of the small clusters, merging them into larger ones, compared to other methods. Infomap and *MCL*_1.5_ had slightly coarser-grained clusters that the ground truth. *MCL*_3_ and *MCL*_5_ often over-clustered and returned finer-grained clusters.

We are aware that the resolution limit helps with Louvain’s performance in experiments involving large clusters; where not searching for smaller clusters helps with the recovery of the large ones [24]. This phenomenon is called field-of-view limit [45] and is known to hinder Infomap’s performance [24]. For local IBD graphs however, field-of-view limit does not pose a challenge as the communities are clique-based and their size are orders of magnitude smaller compare to the total number of nodes. In the PAGE study dataset with 52,000 nodes, the largest cluster we extracted had 726 members.

#### The Effects Of False-Positive Edges

Our experiments show that the supremacy of the Infomap, *MCL*_1.5_, and *MCL*_2_ performances over other methods is stable for false-positive rates ranging from 5% to 50% of the total number of edges. Figure 7 illustrates the effects of false-positives on the performance of algorithms in every metric. High rates of false-positive edges were simulated to simplify detection and comparison of performance patterns. They do not happen in our real data experiments regularly since iLASH, our IBD estimation algorithm, has a low false-positive rate [8]. The statistical power of Infomap and *MCL*_1.5_ stays stable as the number of false-positives grows (Figure 7 E). The power of *MCL*_2_ slightly decreases as the rate of false-positives is increased above 30%. However, it still stays above 0.9. The three methods always recover the disease information simulated for larger clusters, suggesting that they do not: (1) break these clusters into smaller ones, and (2) mix them together as a results of their false-positive connections to each other. This is not true for other clustering methods as their power decreases with higher rates of false-positives, seemingly converging to a minimum value.

The minimum value for the scores is determined by the large clusters that are less structurally affected by the higher rates of false-positive edges. In case of modularity optimizing methods, the lower bound is affected by the resolution limit, as mentioned in the last section. Increasing the number of edges in the graph (by adding false-positive edges), thus has a twofold effect on Louvain and Leiden. First, it increases the chances that two clusters are merged by the methods, especially if their aggregated number of edges is smaller than the resolution limit threshold. Second, it increases the resolution limit itself, intensifying the adverse effects of resolution limit. Figure 10B illustrates how resolution limit threshold increases as the number of false-positives grows. Compared to the resolution limit trends when the number of clusters was increased, increasing false-positives had a subdued effect on the resolution limit. In Figure 10B the percentage of clusters that are larger than the threshold does not approach 0%, explaining the lower bound on statistical power.

AMI score trends slightly differ from power, primarily due to a more pronounced effect of smaller clusters. *MCL*_1.5_ and Infomap yield less stable results. While *MCL*_3_ and *MCL*_5_ have a similar performance to the top performing methods with a false-positive rate of 5%, their performance declines with higher intensity, resulting in the same pattern as their power score. However, due to the lack of baseline adjustment, these two methods continue having the same NMI score as the top performing methods solely by generating a large number of fine-grained clusters (see Figure 12).

*MCL*_5_ and *MCL*_3_ achieved the highest purity scores (Figure 7 C) as purity is biased towards more granular clustering results and can be maximized readily by having each node assigned to a single dedicated cluster. Higher inflation rates result in a fine-grained clusters in MCL, helping the method score higher in purity.

#### False-Negatives Edges

As shown in Figure 8E, the effects of false-negative edges on the power of the algorithms is less pronounced than that of the false-positives edges. While false-negative edges have an adverse effect on the power of MCL and Infomap, they do not affect the power of Louvain and Leiden significantly. Still, even the lowest power scores of *MCL*_1.5_, *MCL*_2_, and Infomap, at a false-negative rate of 50%, is 70% higher than the scores of Louvain and Leiden.

Resolution limit works slightly in favor of Louvain and Leiden here. As discussed in the previous section, resolution limit depends on the number of edges. Higher rates of false-negatives, thus, improve the performances of Leiden and Louvain as they dampen the effects of the resolution limit via lowering the number of edges. Figure 10C displays the effects of increasing the percentage of false-negative edges on the resolution limit. As shown in Figure 8D, the effects of false-negative edges on modularity of the graph are also eviden in the modularity score. While their power score decreased, the top performing algorithms gained higher modularity scores. This is the opposite of what happened when the number of false-positives edges grew; causing modularity to have a higher correlation with power and AMI. Thus, compared to false-positive trends, modularity optimization methods receive a performance boost in terms of both power and AMI scores.

#### Runtimes

Figure 13 displays the average amount of time (in seconds) each method took in our experiments to analyze a dataset as the number of clusters in the dataset grew. The runtime for all methods seem to grow quadratically with respect to the number of simulated clusters. Louvain and Leiden were the fastest methods, analyzing datasets with 5,000 clusters in 0.9 and 0.6 of a second, respectively. The slowest algorithm, Infomap, took 191 seconds on average for the same number of clusters, while the other top performing methods in terms of power, *MCL*_1.5_ and *MCL*_2_, took 30 and 15 seconds on average, respectively. We exclude HCS here since it does not pass our criteria for scalability.

**Figure 13.**
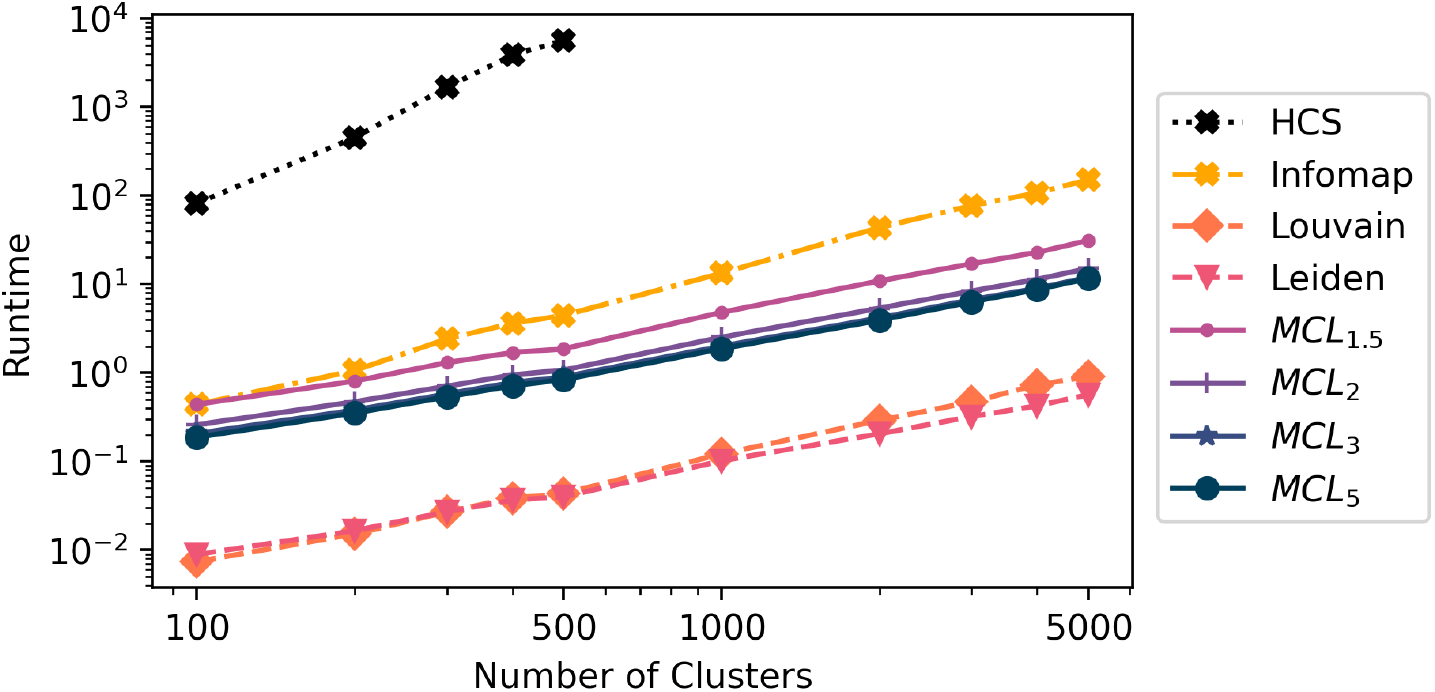
Clustering methods runtimes. Average runtime of the clustering methods for graphs with cluster counts ranging from 100 to 5000 clusters. Each repeated 150 times with various false-positive false-negative ratios ranging from 5% to 50%. HCS algorithm time complexity grows quadratically. It took more than an hour to analyze any graphs with more than 300 clusters. The runtime trends of other methods suggest that they could scale up to analyze Biobank-scale local IBD graphs with tens of thousands of clusters.

#### Highly Connected Subgraphs

DASH, a method that uses HCS, has been a standard tool for IBD mapping in recent years [4]. DASH uses a number of heuristics in its implementation of HCS, which were fine-tuned based on the performance of GERMLINE [46] (an IBD estimation algorithm) in terms of its expected false-positive and false-negative rates in estimating IBD segments. Our experiments, however, cover a range of false-positive/false-negative rates; we do not provide *a priori* knowledge about these parameters to the algorithm in order to have a fair comparison. Moreover, HCS, the oldest clustering method among the five methods we analyzed, does not scale to the size of our experiments. We ran HCS, and the other four algorithms, on a set of 750 small graphs, with cluster counts ranging from 100 to 500. While other algorithms took less than half a second on average to analyze graphs with 100 clusters, HCS took 81.6. This number grew quadratically to 5595 seconds to analyze graphs with 500 clusters (Figure 13). For the same number of clusters, *MCL*_2_ analysis took only 1 second on average.

While we were not able to analyze HCS performance in our simulation and real data analysis due to its scalability issues, our simulations of smaller datasets showed that HCS has a lower statistical power compared to that of Infomap and MCL. The average statistical power of HCS algorithm in these experiments was 0.23 while the top performing algorithm, Infomap had an average score of 0.92.

### Performance on Real Data

We next used the PAGE study dataset to compare the algorithms on real data. First, we ran iLASH over the chromosome 1 genotype data to estimate IBD^[1]^. While false-negative and false-positive edges occur in local IBD graphs due to a variety of phenomena (minimum length of IBD, genotyping errors, phasing errors), our previous analysis suggests iLASH introduces negligible rates of false-positives and false-negatives [8], which prevents high false-positive/false-negative rates in local IBD graphs. We then divided the first chromosome into a set of windows based on the ends of the IBD segments, such that no segments would start or end inside a window (see Figure 1). We generated a local IBD graph for each window, where nodes pertaining to samples that share a segment IBD in that window are connected via an edge. Out of the resulting 8,447 local IBD graphs, we randomly chose 800 (∼ 10%) to cluster using every algorithm. We then calculated the feature-based metric scores of the results. For each local IBD graph, we also generated and analyzed two population specific graphs for African American and Puerto Rican subpopulations, where all the nodes that do not belong to these populations were removed from the graphs.

The real dataset results further demonstrate the effects of the resolution limit on Louvain and Leiden (Table 4). In every population, the two algorithms returned the lowest percentages of node connectivity and highly-connected subgraphs among the clustering methods. Their high rate of edge coverage suggests they are not able to detect false-positive edges. An inflated percentage of missing intra-cluster edges further proves this. Their total clustering of the PAGE data on chromosome 1 requires 43% additional edges in order to turn all the clusters to cliques, compared to *MCL*_1.5_ (top performing method in the simulations) which requires 10% less edges. *MCL*_5_ requires only 19.7% additional edges to achieve the same task, 24% less than Louvain and Leiden.

The score gap between Infomap, *MCL*_1.5_, and *MCL*_2_ on feature-based clustering metrics decreases in the real datasets compared to the simulated ones. This can be partly explained by a lower false-positive rate demonstrated in the high coverage scores achieved by all the methods. To explore this, we trained a linear regressor based on the feature-based metric scores of all algorithms in our simulations to predict false-positive and false-negative rates of the graphs. The linear regressor could predict false-positive and false-negative rates in our simulated graphs with an average error of 2% (std=1%) and 1%(std=2%), respectively. We employed cross validation leaving 20% of the data for testing each round. Using the linear regression model, we estimated that, in our PAGE dataset, the false-positive rate is 2% (std*<* 1%), and the false-negative rate is 24% (std=3%). Similarities between the coverage scores and false-positive rates in our simulations, listed in Table 2, further back up the predicted false-positive rates. For example, the coverage of *MCL*_2_ goes from 96% to 84% as the number of false-positives is increased from 5% to 20%. Based on the coverage scores of *MCL*_2_ on the PAGE dataset (Table 4), we can infer the average false-positive rate of the dataset is under 5%. With fixed values for the false-positive rate, the missing intra-cluster edge rate, listed in Table 3, scores can now be observed to verify our predicted false-negative rates in the same manner.

**Table 2.** Average Coverage scores across all simulated experiments with false-positive rates of 5% and 20%. Higher connectivity scores observed in the real datasets, suggest that the false-positive rates of our real datasets less than 5%.

| | False-Positives | Infomap | Louvain | Leiden | $MCL_{1.5}$ | $MCL_2$ | $MCL_3$ | $MCL_5$ |
| --- | --- | --- | --- | --- | --- | --- | --- | --- |
| Coverage | 5% | 97.33% | 98.82% | 98.83% | 96.86% | 95.80% | 90.24% | 62.90% |
|  | 20% | 87.05% | 91.68% | 91.69% | 86.51% | 83.59% | 67.77% | 28.90% |

**Table 3.**
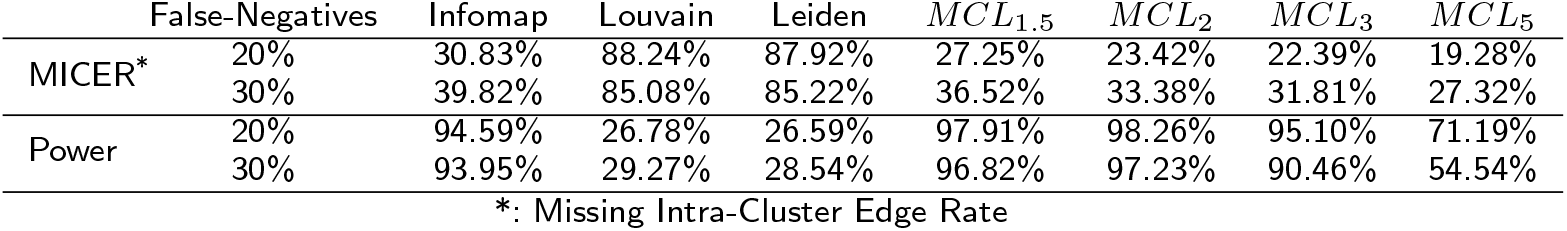
Average missing intra-cluster edge ratio (MICER) scores across all simulated experiments with false-negative rates of 20% and 30%, and false-positive rate of 5%.

| | False-Negatives | Infomap | Louvain | Leiden | $MCL_{1.5}$ | $MCL_2$ | $MCL_3$ | $MCL_5$ |
| --- | --- | --- | --- | --- | --- | --- | --- | --- |
| MICER* | 20% | 30.83% | 88.24% | 87.92% | 27.25% | 23.42% | 22.39% | 19.28% |
|  | 30% | 39.82% | 85.08% | 85.22% | 36.52% | 33.38% | 31.81% | 27.32% |
| Power | 20% | 94.59% | 26.78% | 26.59% | 97.91% | 98.26% | 95.10% | 71.19% |
|  | 30% | 93.95% | 29.27% | 28.54% | 96.82% | 97.23% | 90.46% | 54.54% |

\*: Missing Intra-Cluster Edge Rate

**Table 4.**
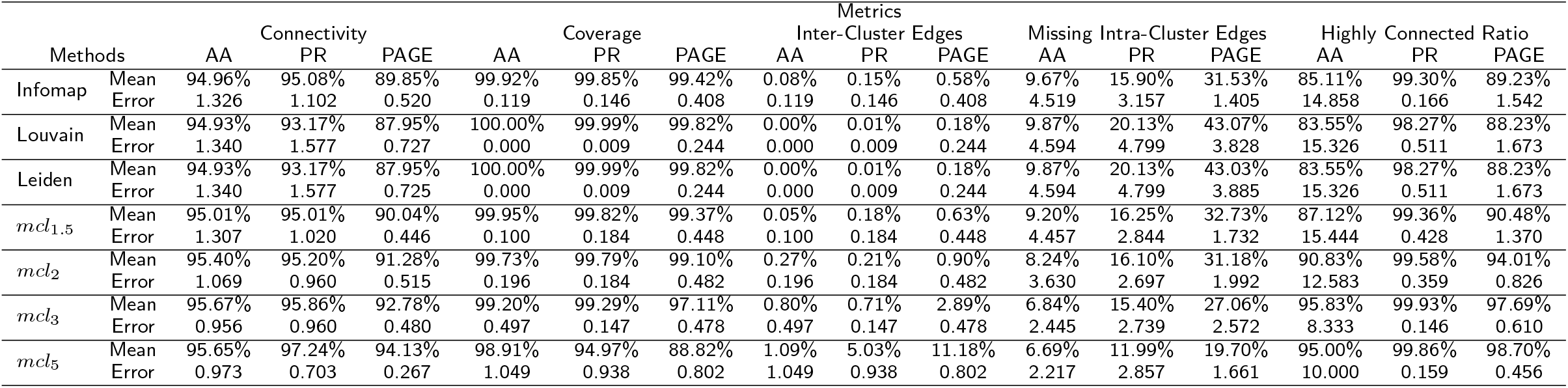
Clustering Algorithm Performance On The PAGE Study Dataset

Focusing exclusively on the simulated graphs with false-positive and false-negative rates close to the ones we estimated for the PAGE study dataset shows a clear superiority for *MCL*_2_ in terms of statistical power. Figure 14 shows the distribution of statistical power scores of all algorithms on simulated graphs with false-positive rate of 5%, and false-negative rates of 20% and 30%. To expand on these results, we simulated 100 graphs, each containing 11,000 clusters (the average number of clusters in a PAGE study dataset local IBD graph) and with realistic false-positive/false-negative rates we estimated. In these simulations, *MCL*_2_ yielded the highest average statistical power score of 98.8%, followed by *MCL*_1.5_ (98.6%), *MCL*_3_ (97.6%) and Infomap (95.5%). Louvain and Leiden had the lowest score at 35%, considerably lower than the *MCL* methods.

**Figure 14.**
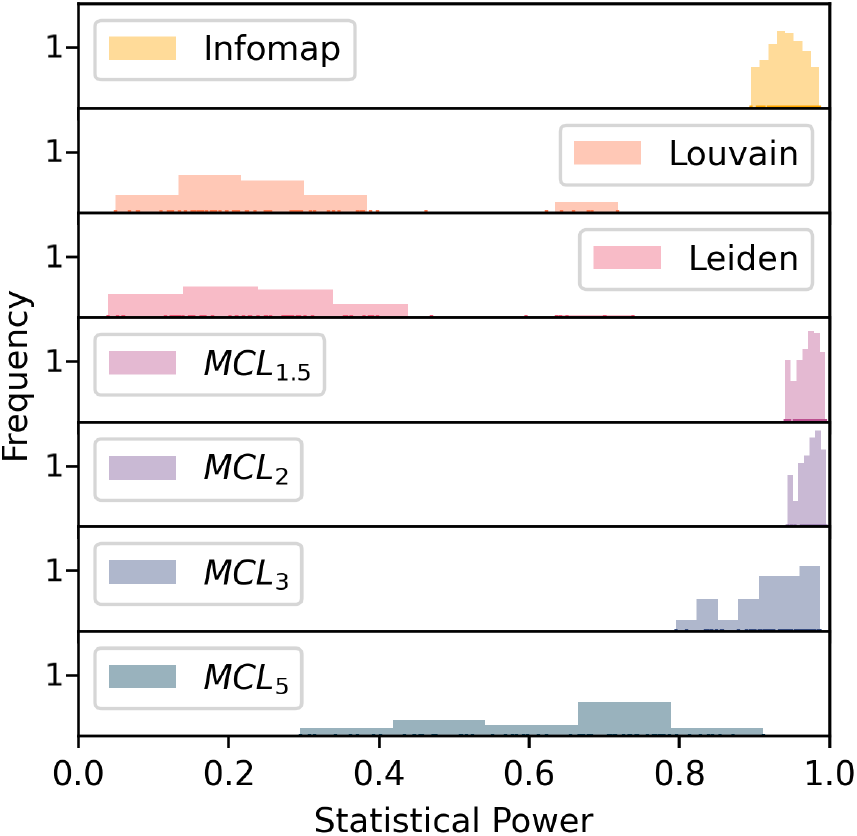
Distribution of Statistical Power Scores. The distribution of power scores in 60 experiments with a false-positive rate of 5% and false-negative rate of 20% and 30%. These rates were closest to our predicted rates in the PAGE dataset.

## Discussion

We proposed a realistic approach to simulate local IBD graphs that addresses distinctive properties of such graphs, and analyzed the suitability of scalable clustering algorithms for IBD Mapping. Our measurements and analysis found that common benchmark graphs such as LFR [23] are not suited to represent local IBD graphs due to the aforementioned properties. Our benchmark, thus, provided us with a ground truth for analyzing a group of scalable clustering algorithms and common clustering metrics for the purpose of local IBD clustering for the first time.

We demonstrated that available analyses on clustering algorithms and clustering metrics do not apply to local IBD graphs, further stressing the importance of our analysis. Comparing feature-based clustering metrics, including modularity, against AMI and statistical power showed that metrics based on community structure cannot be considered sufficient substitutes for statistical power in IBD mapping. We also found out that even metrics that are based on ground truth, specifically AMI, are not a perfect substitute for statistical power despite having higher correlation with it when compared to feature-based metrics. Errors in small clusters (which are ignored in IBD mapping) affect AMI more than power, causing discrepancies.

While feature-based clustering metrics cannot predict statistical power, we showed that they can help with realistic dataset specific simulations of local IBD graphs. The simulations determine the fittest clustering algorithm in terms of statistical power.

As suggested by Emmons et al [24], the definition and structure of communities under study should derive the decision on what clustering methods to use. Our real dataset analysis shows various population structures may also require specific clustering approaches. *MCL*_2_ generally performed better than the other methods in our realistic experiments. We believe *MCL*_2_ to be the best clustering method for general IBD mapping purposes. However, various datasets and IBD estimation algorithms have different false-positive/false-negative rates; necessitating dataset specific simulations in order to find the fittest clustering algorithm. A preliminary analysis using *MCL*_2_ to calculate its missing intra-cluster edges and coverage scores on the data can help estimate realistic false-positive/false-negative rates for such simulations.

We showed that the cluster size distribution of IBD graphs, which is heavily skewed towards smaller clusters, could result in numerous groups of small clusters being aggregated by clustering methods, specially for methods that are based on greedy modularity optimization. Moreover, we found further evidence that the performance of greedy modularity optimizing methods is dependent on the size of the graph being analyzed, making them unpredictable.

Novel clustering methods, such as spectral clustering [47] and neural network clustering [48], do not scale to the size of the graphs in IBD Mapping. Methods based on spectral decomposition of adjacency matrix took significantly longer to process even smallest graphs in our experiments. (Note that a local IBD clustering process involves clustering thousands of graph datasets independently). Moreover, finding the optimal cluster count from the decomposition, or choosing a numerical clustering method for the eigenvectors introduces optimization problems that have a different scope than that of this paper. This, together with the exaggerated gap in run time resulted in the exclusion of spectral clustering methods from our analysis. Also, while spectral clustering can help with modularity optimization [47], we have demonstrated that optimizing modularity should not be a priority for local IBD clustering.

Methods based on neural networks were three orders of magnitude slower than other methods (4000-fold increase in runtime). One of the main strengths of neural network based methods such as node-2-vec is their ability to incorporate knowledge from large data sets and apply it on other data sets [48]. In our case, however, we are working with a large number of smaller graphs, which we aim to analyze independently. Moreover, the extreme disjointedness of our graphs is unfavorable to the both neural network based and spectral based approaches [48, 47].

While IBD mapping can help us understand the genetic origins of some traits, its potential is bound by the capabilities of its clustering approach. Even slight clustering errors can negatively affect the accuracy due to the small size of the local IBD communities. Clustering algorithms such as *MCL* can help alleviate these effects by better eliminating erroneous IBD data. We plan to utilize the *MCL* algorithm to conduct a large IBD mapping analysis on the UK Biobank dataset.

We believe distinctive properties of UK Biobank, such as its size, and health record availability, together with power of IBD mapping will help us find novel genetic associations.

We plan to add two functionalities to our benchmark algorithm. First, we aim to design a realistic approach to simulate edges weights for the graphs that represent IBD segments length. Augmenting local IBD graphs with segment lengths as edge weights can help clustering methods (that support weights) detect false-positives more accurately. The longer the segment, the lower the probability of it being a false-positive edge. Second, we plan to simulate overlapping local IBD graphs, where a group of IBD graphs are merged and processed together to save computing resources. In order to reduce the number local IBD graphs to process, we can aggregate them in groups via dividing the chromosome into windows of static length (for example 0.5 cM). We aim to evaluate clustering algorithms’ power in detecting overlapping communities in our benchmark. Simulating these two phenomena requires a genetic coalescence simulation that is outside the scope of this paper.

## Conclusion

We demonstrated the shortcomings of LFR, the common benchmark algorithm, in simulating local IBD graphs, and proposed a realistic benchmark that addresses them. We evaluated the performances of four modern clustering algorithms in the context of local IBD graphs and IBD mapping and compared them with the standard algorithm, HCS, in clustering 3,374,500 simulated and ∼8,800,000 real clusters. We showed that utilizing clustering approaches that do not fit the target dataset properties results in poor statistical power of the downstream analyses, such as IBD mapping. We introduced an approach that uses our benchmark to find the fittest clustering algorithm for a dataset. It can also be utilized to optimize clustering algorithm parameters. We found MCL_2_ as the most appropriate algorithm for our purpose and dataset. Furthermore, our findings show that greedy modularity optimization approaches are not appropriate for local IBD graph clustering in general. These findings will make IBD mapping on large scale datasets such as the UK Biobank possible, paving the way for the efficient discovery of novel causal rare variants.

## Declarations

### Availability of data and materials

The code package used to generate the benchmark, clustering local IBD graphs, and calculating the scores is available on our Github repository. To calculate *R*^2^ scores, we use scikit Python library [44].

The simulated data generated and analyzed in this study is publicly available on Mendeley Data (DOI: 10.17632/72sh3pp4m2.1). It can also be regenerated using the our benchmark code on Github using the parameters mentioned in the results section.

The real dataset was generated using the PAGE study IBD segment data, generated as described in [8]. The Population Architecture using Genomics and Epidemiology (PAGE) study data is available through dbgap at accession number phs000356. IBD segments of 51,520 individuals were used to generate the local IBD graphs using the scripts available on the Github repository.

### Author’s contributions

J.L.A, C.R.G, and G.M.B led the development of the approach. R.S, J.L.A, and C.R.G designed the benchmark with technical input from G.M.B, K.B, and E.E.K. R.S. implemented the pipeline and ran the simulations with input on clustering algorithms and metrics from K.L. R.S. and K.B chose the metrics and analyzed the simulation results with input from J.L.A and C.R.G. R.S. ran the clustering analyses on the PAGE dataset with input from G.M.B, E.E.K and C.R.G. R.S. and J.L.A wrote the manuscript with critical edits from all authors. All authors provided revisions and edits to the submitted manuscript.

## 1 Acknowledgements

Not Applicable

## 2 Ethics approval and consent to participate

Not Applicable

## Footnotes

[1] We chose chromosome 1 since it was the largest chromosome without any regions of low complexity in the PAGE dataset.

